# Neurodegenerative and neurodevelopmental roles for bulk lipid transporters *VPS13A* and *BLTP2* in movement disorders

**DOI:** 10.1101/2024.12.30.630795

**Authors:** Sarah D. Neuman, Rajan S. Thakur, Scott J. Gratz, Kate M. O’Connor-Giles, Arash Bashirullah

## Abstract

**Background:** Bridge-like lipid transfer proteins (BLTPs) mediate bulk lipid transport at membrane contact sites. Mutations in BLTPs are linked to both early-onset neurodevelopmental and later-onset neurodegenerative diseases, including movement disorders. The tissue specificity and temporal requirements of BLTPs in disease pathogenesis remain poorly understood.

**Objectives:** To determine the age-of-onset and tissue-specific roles of *VPS13A* and *BLTP2* in movement disorder pathogenesis using *Drosophila* models.

**Methods:** We generated tissue-specific knockdowns of the *VPS13A* ortholog (*Vps13*) and the *BLTP2* ortholog (*hobbit*) in neurons and muscles of *Drosophila*. We analyzed age-dependent locomotor behavior, neurodegeneration, and synapse development and function.

**Results:** Neuron-specific loss of the *VPS13A* ortholog caused neurodegeneration followed by age-onset movement deficits and reduced lifespan, while muscle-specific loss affected only lifespan, revealing neurodegeneration and myopathy as independent comorbidities in *VPS13A* disease. In contrast, neuronal loss of the *BLTP2* ortholog resulted in severe early-onset locomotor defects without neurodegeneration, while muscle loss impaired synaptogenesis and neurotransmission at the neuromuscular junction (NMJ).

**Conclusions:** *VPS13A* maintains neuronal survival, while *BLTP2* orchestrates synaptic development. *VPS13A* function in muscle does not play a role in movement defects. The phenotypic specificity of BLTP function provides mechanistic insights into distinct disease trajectories for BLTP-associated movement disorders.

## INTRODUCTION

Loss-of-function mutations in *VPS13A* are associated with the rare movement disorder chorea-acanthocytosis, now known as *VPS13A* disease (1). This condition is inherited in an autosomal recessive manner and usually manifests in adulthood (mean age-of-onset 30 years) (1). Patients with *VPS13A* disease exhibit a spectrum of neurologic symptoms, including involuntary and unpredictable muscle movements (chorea), tics, abnormal muscle contractions (dystonia), and behavioral and cognitive changes (1). This condition is progressive, leading to significant disability and premature death. Currently, there is no cure for *VPS13A* disease, and treatment centers around managing symptoms to improve patient mobility and comfort. Traditionally, *VPS13A* disease has been thought to be primarily neurodegenerative in nature; however, some studies suggest that myopathy may also be a key player in disease etiology (2–4). Specifically, a recent study obtained muscle biopsies from patients with *VPS13A* disease and found significant variation in muscle fiber size and organization (4), pointing to myopathy as a likely contributor to disease. The relative roles of *VPS13A* dysfunction in neurons vs. muscle in disease pathology remain unclear.

VPS13A is a member of the recently defined bridge-like lipid transfer protein (BLTP) superfamily; these large proteins localize to membrane contact sites and mediate bulk transfer of lipids between organelle membranes (5–20). There are five basic members of the BLTP superfamily, several of which have two or more paralogs in humans: *BLTP1*, *BLTP2*, *BLTP3A/B*, *ATG2A/B* (*BLTP4A/B*), and *VPS13A-D* (*BLTP5A-D*) (21). All these proteins share a characteristic bridge-like shape, and all contain an interior hydrophobic cavity allowing them to function like lipid superhighways, potentially promoting an uninterrupted flow of lipids between organelle membranes (10). Evidence so far suggests that BLTPs are agnostic to the lipids they transport, and specificity, if any, is likely mediated by partner proteins, such as scramblases that help equilibrate the inner and outer leaflets of membranes during lipid transfer (8,22–25). In fact, mutations in *XK*, a scramblase partner of *VPS13A* (22,26), are associated with McLeod syndrome, a neuroacanthocytosis disorder with clinical manifestations similar to *VPS13A* disease (27). Specificity in BLTP function is also directed by partner proteins that promote localization to specific membrane contact sites; for example, VPS13A interacts with adapter proteins that promote its localization to endoplasmic reticulum (ER)-mitochondria and ER-lipid droplet contact sites (5,28). Ongoing work is focused on understanding the mechanisms that drive BLTP localization and function at membrane contact sites.

Like *VPS13A*, mutations in other members of the BLTP superfamily are associated with neurodegenerative and neurodevelopmental diseases (reviewed in (10,11)). Most recently, *de novo* variants in *BLTP2* were associated with neurodevelopmental disorders (29) and conditions like autism (30,31) and schizophrenia (32). Development of the nervous system requires proper control of neuronal connectivity, mediated by neurogenesis, synaptogenesis, and neuronal pruning, among other processes (33). We have previously characterized the *Drosophila* ortholog of *BLTP2*, named *hobbit*, and found that loss-of-function mutations in *hobbit* result in animals with a dramatic small body size and post-embryonic lethality (34). The small body size is caused by secretion defects within the neuroendocrine axis that controls insulin/insulin-like growth factor signaling (IIS), a central regulator of both metabolism and systemic growth in all animals (35–37). Work by us and others has shown that BLTP2/Hobbit localizes to ER-plasma membrane (PM) contact sites (18,20,38). Although the function of *BLTP2/hobbit* in body size control has been characterized, much remains unknown about the function of this protein in neurons and neurodevelopmental disorders.

In this study, we use *Drosophila* as a genetically tractable model system to dissect age-of-onset and tissue-specific roles for two BLTPs, *VPS13A* and *BLTP2*, which have clinical roles in neurodegeneration and neurodevelopment, respectively. All BLTPs are highly conserved; the *Drosophila* genome contains an ortholog of *BLTP2* (*hobbit*) and three paralogs of *VPS13* (*Vps13, Vps13B,* and *Vps13D*). We find that loss of *VPS13A/Vps13* in neurons causes neurodegeneration followed by age-onset movement defects and reduced lifespan, while loss of this protein in muscle causes reduced lifespan without movement defects or neurodegenerative changes. Thus, our results suggest that movement disorders characteristic of *VPS13A* disease may be caused primarily by neurodegeneration, not myopathy; however, myopathy is likely an independent comorbidity in *VPS13A* disease. Additionally, we find that loss of *BLTP2/hobbit* in neurons causes early-onset movement defects without substantial neurodegenerative changes; instead, we show that this protein is required postsynaptically for proper motor synapse formation during development. This study describes the first neurodevelopmental function of *BLTP2*. Overall, our results show that BLTPs have distinct tissue-specific functions in neurodegenerative and neurodevelopmental movement disorders.

## METHODS

### *Drosophila* stocks and husbandry

Stocks were obtained from the Bloomington *Drosophila* Stock Center: *w^1118^* (RRID:BDSC_3605), *elav-GAL4* (RRID:BDSC_458), *24B-GAL4* (RRID:BDSC_1767), *act-GAL4* (RRID:BDSC_3954), *UAS-*

*luciferase* (RRID:BDSC_35788), *UAS-Vps13-RNAi* (RRID: BDSC_38270). We used FlyBase (release FB2024_05) (39) to find stock information. *hob^2^, hob^3^*, and *UAS-hobbit-RNAi 1* were previously described and validated (34). To generate *endogenous hobbit-GFP*, we obtained a fosmid containing C-terminal superfolder GFP (sGFP)-tagged *hobbit* and surrounding genomic sequences (*Drosophila* TransgeneOme; (40)); schematic in Fig. S1. We sequence-verified the sGFP tag and injected the fosmid into *VK00027* flies for phiC31-mediated site-directed integration using standard methods (Rainbow Transgenic Flies, Inc.). All *Drosophila* crosses were grown on standard cornmeal molasses media in density-controlled bottles or vials kept in a 25°C incubator with a 12 h light-dark cycle. Pupa imaging and body size and lethal phase analysis were performed as previously described (34).

#### Adult climbing and larval crawling assays

Adult climbing assays were performed as described in (41). Climbing assays on comparably aged animals were conducted in parallel; 3-4-day-old *elav>luc*, *elav>Vps13(i)*, and *elav>hob(i)* were all analyzed together. Thus, the same control (*elav>luc*) data is used in Figs. 1D and 3B. Larval crawling analysis was performed following published protocols (42) and analyzed using FIJI (RRID: SCR_003070) (43,44). See extended methods for additional details.

**Figure 1.**
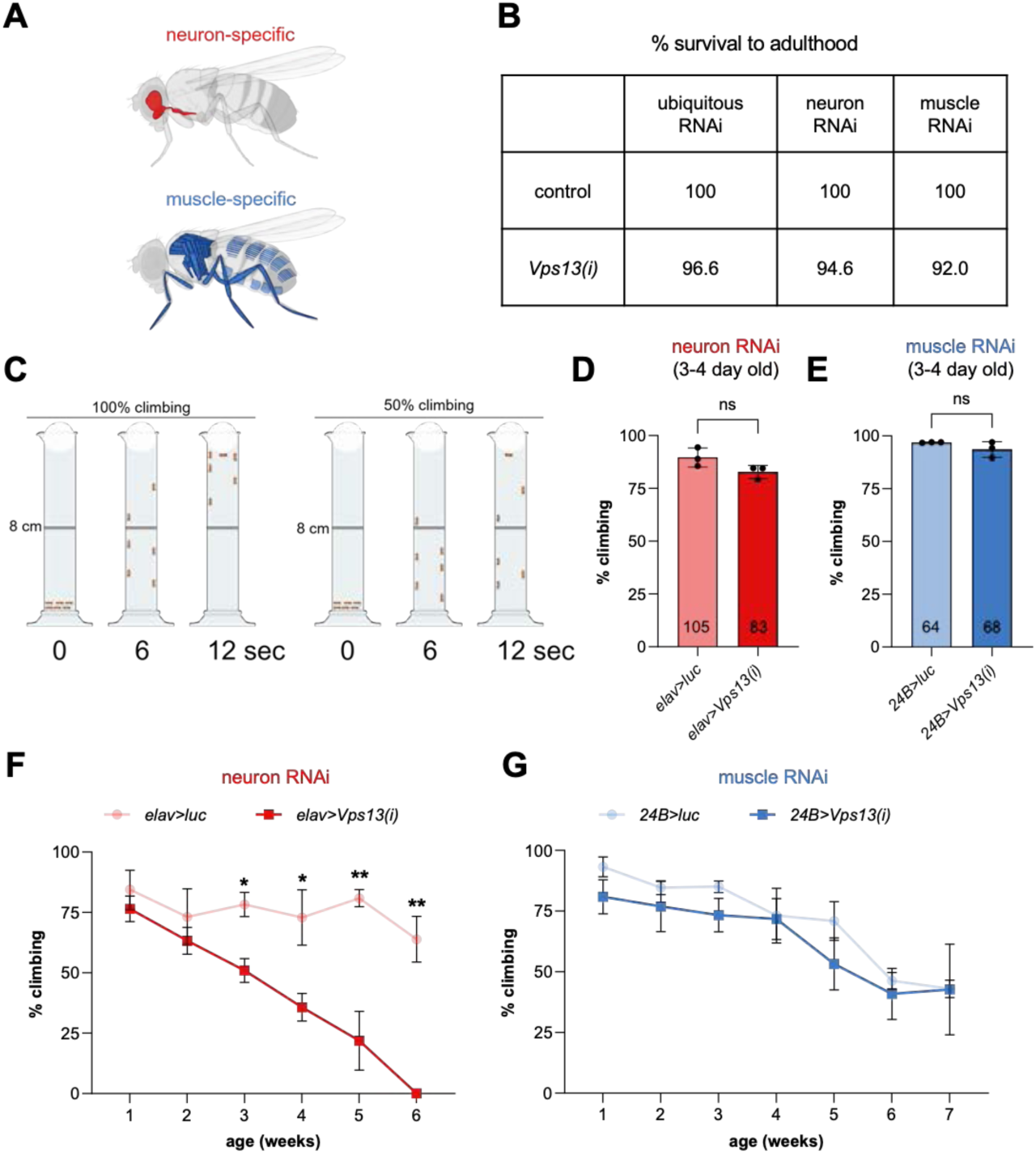
Loss of *Vps13* in neurons results in age-onset movement defects. **(A)** Schematic of tissue-specific knockdown in *Drosophila*. Specific knockdown in neurons (red) was done using the pan-neuronal driver *elav-GAL4* . Specific knockdown in muscles (blue) was done using the pan-muscle driver *24B-GAL4*. **(B)** Table depicting percent survival to adulthood upon ubiquitous (*act-GAL4*), neuron-specific (*elav-GAL4*), or muscle-specific (*24B-GAL4*) knockdown of *Vps13* compared to a genotype-matched control (*GAL4* driver with *UAS-luciferase* (*luc*), e.g., *act>luc*). *n*≥50 animals analyzed per genotype. **(C)** Schematic depicting adult fly climbing assays, with examples of 100% climbing ( left) and 50 % c l imbing ( r ight). In experiments, flies were analyzed in groups of *n*∼10. **(D-E)** Climbing assays in 3–4-day old neuron-specific **(D)** or muscle-specific **(E)** *Vps13* knockdown flies and paired controls. Total *n* shown on each bar. For each genotype, flies were collected from three independent genetic crosses; individual data points represent average climbing for each of these biological replicates. **(F-G)** Climbing assays in control (round symbols) and neuron-specific knockdown, red **(F)**, or muscle-specific knockdown, blue **(G)**, of *Vps13* (square symbols) in aging flies. *elav>luc n*=79; *elav>Vps13(i) n*=68; *24B>luc n*=75; *24B>Vps13(i) n*=70. All samples analyzed in biological triplicate. Graphs show mean±S.D. Significance calculated using unpaired, two-tailed *t*-tests. \**p*<0.05; \*\**p*<0.01; ns=not significant.

### Histology and neurodegeneration index (NI) quantification

Fly heads were prepared for histology, embedded in paraffin, and sectioned using standard methods (45); see extended methods for details. Images were obtained using a BioTek Lionheart FX Automated Microscope (10x objective) with Gen5 v.3.11 software, then rotated and cropped post-acquisition in Adobe Photoshop CS6. NI were quantified using the criteria described in (45). The number and location of vacuoles in the brain determined the NI: 0 = no vacuoles; 1 = a few, small vacuoles mainly in optic lobes in only a few sections; 2 = many vacuoles in many sections, mainly in optic lobes, but possibly some in central brain; 3 = vacuoles become prominent in central brain and are abundant in optic lobes; 4 = many vacuoles in central brain, and some vacuoles in optic lobes and central brain are large. NI scoring was performed blind to age and genotype. Comparably aged samples were collected and analyzed in parallel; thus, the same 1-2-day-old and 1-week-old NI data is used in Figs. 2D and 3E.

**Figure 2.**
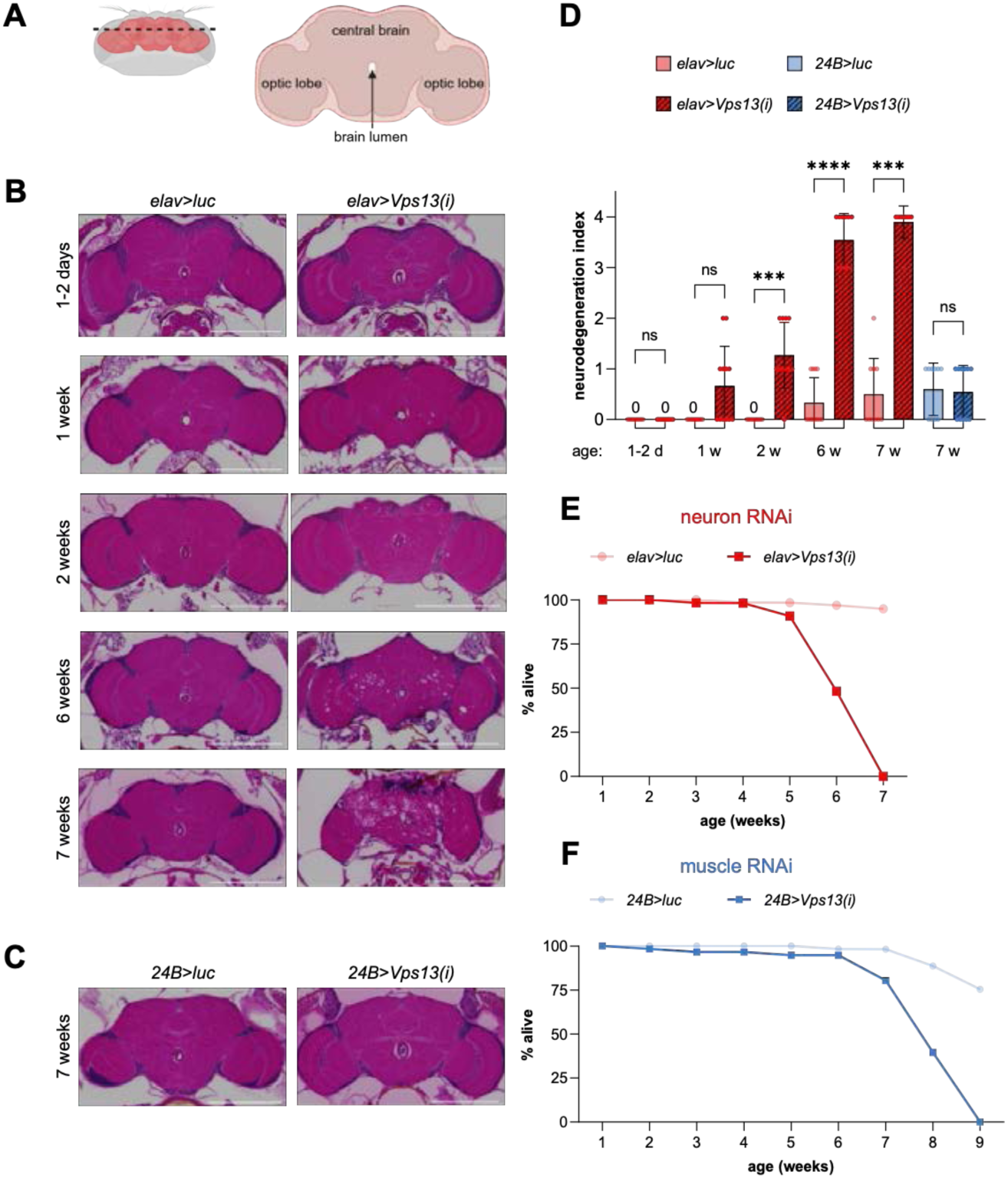
Neuron-specific loss of *Vps13* results in age-dependent neuro-degeneration and shortened lifespan, while muscle-specific loss results in only shortened lifespan. **(A)** Schematic of the adult *Drosophila* brain. For histology, serial sections spanning the entire head (from anterior to posterior) were obtained. Major brain regions visible in histological sections are labeled. **(B)** Representative hematoxylin and eosin (H&E) stained sections of adult fly brains (midbrain region) from control (*elav>luc*) and neuron-specific *Vps13* knockdown (*elav>Vps13(i)*) at the indicated ages. Scale bars 200 µm. **(C)** H&E-stained sections of control (*24B>luc*) and muscle-specific *Vps13* knockdown (*24B>Vps13(i)*) brains from 7-week-old flies. Scale bars 200 µm. **(D)** Quantification of neurodegeneration index for samples shown in panels B and C. See Methods section for criteria used to define each index. *n≥*10 brains analyzed for each genotype/age. **(E-F)** Survival curves for neuron-specific, red **(E),** or muscle-specific, blue **(F)**, control (round symbols) and *Vps13* knockdown (square symbols) flies. Note that the same group of flies was used for both the climbing assays depicted in Fig. 1F-G and the lifespan analysis shown h e r e . G r a p h s s h o w m e a n ± S . D . Significance calculated by Kruskal-Wallis followed by Dunn’s multiple comparison test. \*\*\**p*<0.001; \*\*\*\**p*<0.0001; ns=not significant.

**Figure 3.**
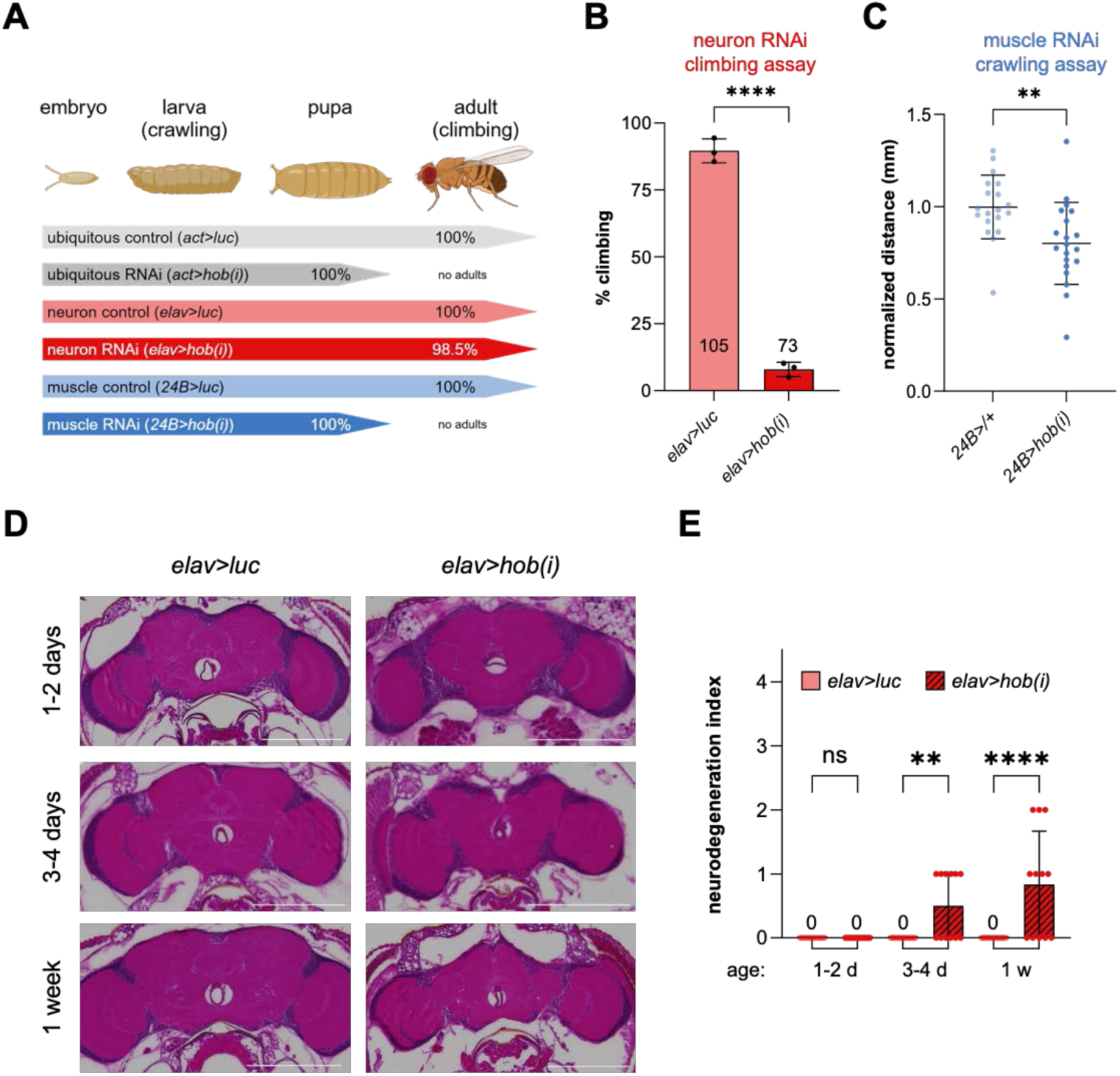
Neuron- and muscle-specific loss of *hobbit* impairs locomotion without neurodegeneration. **(A)** Summary of the fly life cycle; note that locomotion can be assessed in larvae using crawling assays and in adults using climbing assays. Animals with ubiquitous or muscle-specific knockdown of *hobbit* perish before adulthood, while nearly all animals with neuron-specific knockdown of *hobbit* survive to adulthood. *n*≥50 animals analyzed per genotype. **(B)** Adult climbing assays in 3-4-day old flies show that neuron-specific knockdown of *hobbit* significantly impairs locomotion. Total *n* shown on bars. For each genotype, flies were collected from three independent genetic crosses; individual data points represent average climbing for each of these biological replicates. Graph shows mean±S.D. Significance calculated by unpaired, two-tailed *t*-test. **(C)** Larval crawling assays show that muscle-specific knockdown of *hobbit* significantly impairs locomotion. Graph shows mean±S.D. with data points representing individual animals. *n≥*19 animals analyzed per genotype. Significance calculated by Mann-Whitney test. **(D)** Representative H&E-stained sections of control (*elav>luc*) and neuron-specific knockdown of *hobbit* (*elav>hob(i)*) brains (midbrain region) at the indicated ages. Scale bars 200 µm. **(E)** Neurodegeneration index quantification of the samples shown in panel D. *n*≥10 brains analyzed per genotype/age. Graph shows mean±S.D. Significance calculated using Kruskal-Wallis followed by Dunn’s multiple comparison test. \*\**p*<0.01; \*\*\*\**p*<0.0001; ns=not significant.

### Larval neuromuscular junction (NMJ) imaging and synaptic bouton quantification

NMJ preparation was performed as previously described (46); details in extended methods. Images were acquired on a Nikon A1R HD confocal microscope with a Plan-Apo 60x 1.49 NA oil-immersion objective. Boutons were quantified in FIJI (RRID: SCR_003070) (43,44) masked to genotype.

### Electrophysiology

Electrophysiology was performed as previously described (47); details in extended methods. At least 30 consecutive excitatory junction potentials (EJPs) were recorded for each cell and analyzed in pClamp to obtain mean amplitude. Quantal content (QC) was calculated as the ratio of mean EJP amplitude to mean mEJP amplitude.

## DATA AVAILABLITY

All relevant data is included in the manuscript.

## RESULTS

### A neuron-specific requirement for *Vps13* in age-onset movement defects

Inactivating mutations in human *VPS13A* are associated with chorea-acanthocytosis, now known as *VPS13A* disease (1,48). While this disease has historically been thought to be primarily neurodegenerative in nature, recent studies suggest that myopathy may also play a critical role in disease etiology (2–4). We used the fruit fly *Drosophila melanogaster* as a model system to separate the functional roles of *VPS13A* in neurons and in muscle (Fig. 1A). Loss-of-function mutants of the fly *VPS13A* ortholog *Vps13* are reported to be viable (49,50); consistent with this result, we found that ubiquitous knockdown of *Vps13* did not substantially affect viability, since >95% of the animals emerged as adults (Fig. 1B). The *Vps13-RNAi* construct we used was previously shown to significantly reduce Vps13 protein levels (50). Similarly, neuron-specific or muscle-specific knockdown of *Vps13* had little effect on viability (Fig. 1B), demonstrating that this gene is dispensable for survival into adulthood. We then examined potential functional consequences of *Vps13* loss in neurons and in muscle using climbing assays, which are a well-accepted method to assess motor function in flies (41,51) (Fig. 1C). Young (3-4-day old) flies with neuron-specific (Fig. 1D) or muscle-specific (Fig. 1E) knockdown of *Vps13* did not display any climbing defects. This began to change as the flies aged, however. Flies with neuron-specific knockdown of *Vps13* did not exhibit any climbing deficits compared to controls at one or two weeks of age (Fig. 1F); however, by three weeks of age, the *Vps13* neuronal knockdown flies exhibited significant climbing defects, which continued to decline until no animals were able to properly climb by six weeks of age (Fig. 1F). In contrast, muscle-specific knockdown of *Vps13* had no significant effect on climbing ability even at seven weeks of age when compared to genotype-matched controls (Fig. 1G). Together, these results show that *Vps13* is required specifically in neurons for proper control of movement during aging.

### Neuronal loss of *Vps13* results in age-onset neurodegeneration and shortened lifespan

Human patients with *VPS13A* disease exhibit signs of neurodegeneration; MRI or CT imaging of living patients frequently shows atrophy of the putamen and caudate nucleus, basal ganglia, and hippocampus (1,52,53). Additionally, histological studies of post-mortem brains show cellular signs of neurodegeneration (54). However, it is difficult to determine if movement disorder symptoms or neurodegeneration appear first during disease progression. To address this question, we used histology to examine the brains of aging flies with neuron-specific loss of *Vps13* for signs of neurodegenerative changes (Fig. 2A). The brains of young (1-2-day old) flies with neuron-specific knockdown of *Vps13* were indistinguishable from controls (Fig. 2B, D). However, by one week of age, some brains from flies with neuronal loss of *Vps13* began to show vacuoles (Fig. 2B, D), and the abundance and size of vacuoles continued to progress at two weeks of age (Fig. 2B, D). By six weeks, the brains of flies with neuron-specific loss of *Vps13* exhibited dramatic neurodegenerative changes, with prominent and frequently large vacuoles present in both the optic lobes and central brain (Fig. 2B, D). Importantly, flies with neuronal loss of *Vps13* did not display movement deficits until three weeks of age (see Fig. 1F), showing that neurogenerative changes preceded the onset of movement disorders. In parallel to assessing the movement capabilities of flies with neuron-specific knockdown of *Vps13*, we assessed lifespan. Interestingly, flies with neuron-specific knockdown of *Vps13* began to die at five weeks of age, with all animals perishing by seven weeks; in contrast, few genotype-matched controls died during this timeframe (Fig. 2E; note the maximum lifespan of flies at 25°C is ∼90 days/∼13 weeks (55)). No significant advancement in neurodegeneration was visible in these deceased flies (Fig. 2B, D). Together, this data shows that neurodegenerative changes precede movement disorders and lethality upon neuron-specific loss of *Vps13*.

### Muscle-specific loss of *Vps13* results in shortened lifespan without neurodegeneration

Muscle-specific knockdown of *Vps13* did not cause climbing defects during aging (see Fig. 1G); therefore, *Vps13* function in muscle does not appear to influence movement disorders. Accordingly, the brains of seven-week-old flies with muscle-specific knockdown of *Vps13* showed only mild neurodegeneration consistent with normal aging seen in genotype-matched controls (Fig. 2C-D). However, surprisingly, flies with muscle-specific knockdown of *Vps13* exhibited a shortened lifespan compared to controls (Fig. 2F). This data shows that loss of *Vps13* in muscle does not affect locomotion but does play an important role in determining animal lifespan. Thus, our data suggests that myopathy is likely an independent and critical comorbidity in patients with *VPS13A* disease.

### Neuron-specific loss of *BLTP2/hobbit* causes early-onset movement defects without neurodegeneration

Clinically, altered neurodevelopment is often associated with movement disorders; for example, autistic individuals frequently experience ataxia and other motor coordination challenges (56–58). A recent preprint identified a novel link between *de novo* variants in human *BLTP2* and neurodevelopmental disorders (29); additionally, *BLTP2* variants have been associated with autism and schizophrenia in large-scale sequencing studies (30–32). Although ubiquitous knockdown of *hobbit* resulted in lethality prior to adulthood (34) (Fig. 3A), surprisingly, neuron-specific knockdown of *hobbit* did not affect survival, since nearly all animals emerged as adults (Fig. 3A). Therefore, given the genetic connection between *BLTP2* and neurodevelopmental disorders, we used climbing assays in young adults to test for locomotion deficits upon neuron-specific loss of *hobbit*. Strikingly, 3–4-day old flies exhibited dramatic climbing defects (Fig. 3B), indicative of severe impairments in movement. In fact, we observed that even newly emerged flies tended to linger abnormally at the bottom of vials, unlike controls, which climbed to the lid of vials. Histological analysis of brains from 1–2-day old flies with neuron-specific knockdown of *hobbit* did not reveal any obvious abnormalities (Fig. 3D-E); however, a few vacuoles were observed in the optic lobes of some brains from 3–4-day old flies (Fig. 3D-E). This phenotype was progressive, since at one week of age, some *hobbit* knockdown brains exhibited vacuolation in both the optic lobes and the central brain (Fig. 3D-E). Although 3-4-day old flies with neuronal knockdown of *hobbit* exhibited climbing defects akin to 5-week-old flies with neuronal knockdown of *Vps13* (*c.f.* Figs. 1F and 3B), the vacuolation of *hobbit* knockdown brains was quite mild compared to *Vps13* knockdown brains (*c.f.* Figs. 2B, D and 3D-E). Thus, while vacuolation of the neuronal *hobbit* knockdown brains could be indicative of premature aging, neurodegeneration does not appear to cause the dramatic movement defects seen in these flies. Unfortunately, the severely impaired locomotion of *hobbit* neuronal knockdown flies made analysis of lifespan difficult, since 1-week-old flies were frequently trapped in the food. Overall, these results show that *BLTP2/hobbit* plays an essential role in neurons, since neuron-specific loss of *hobbit* results in dramatic movement defects in young adult flies.

### *BLTP2/hobbit* is required in muscle for animal survival and normal locomotion

Given that our data showed that *Vps13* is required in muscle for normal lifespan (see Fig. 2F), we also tested for any effects upon muscle-specific loss of *hobbit*. Muscle-specific knockdown of *hobbit* did not produce any adult flies (Fig. 3A); all animals died during pupal development. Without adults, we cannot use climbing assays to test locomotion. However, since animals with muscle-specific loss of *hobbit* survive past larval stages and into pupal development, we used larval crawling assays (Fig. 3A) to test if these animals have any movement deficits. We saw a significant reduction in larval crawling upon muscle-specific knockdown of *hobbit* compared to controls (Fig. 3C). Together, these data show that *hobbit* is essential in muscle for survival to adulthood and for normal locomotion in juvenile stages.

### *BLTP2/hobbit* has a specific postsynaptic role during synaptogenesis

All muscle types (skeletal, cardiac, and smooth) are targets of motor neurons; thus, muscle is the postsynaptic cell at neuromuscular junctions (NMJs) (59). At well-characterized *Drosophila* larval NMJs, individual motor neurons target a single postsynaptic muscle cell, forming a stereotyped number of varicosities called boutons, each containing multiple synapses (Fig. 4A). These synaptic connections are initially established during embryogenesis; then, as muscle fibers increase in size during development, boutons are continuously added to maintain the proper synaptic drive required for locomotion (60). Interestingly, *hobbit* mutant animals exhibited a substantial reduction in boutons at the NMJ (Fig. 4B-C). Importantly, bouton number was still significantly reduced when the smaller muscle area of *hobbit* mutant animals was considered (Fig. 4C). To further validate a role for *hobbit* in determining bouton counts, we generated an endogenously regulated *hobbit-GFP* rescue construct (see Methods and Fig. S1 for details). Synapse formation in *hobbit* mutant animals was fully rescued by one copy of the *endogenous hobbit-GFP* construct (Fig. 4B-C), thereby validating the functional role of *hobbit* in regulating synapse formation.

**Figure 4.**
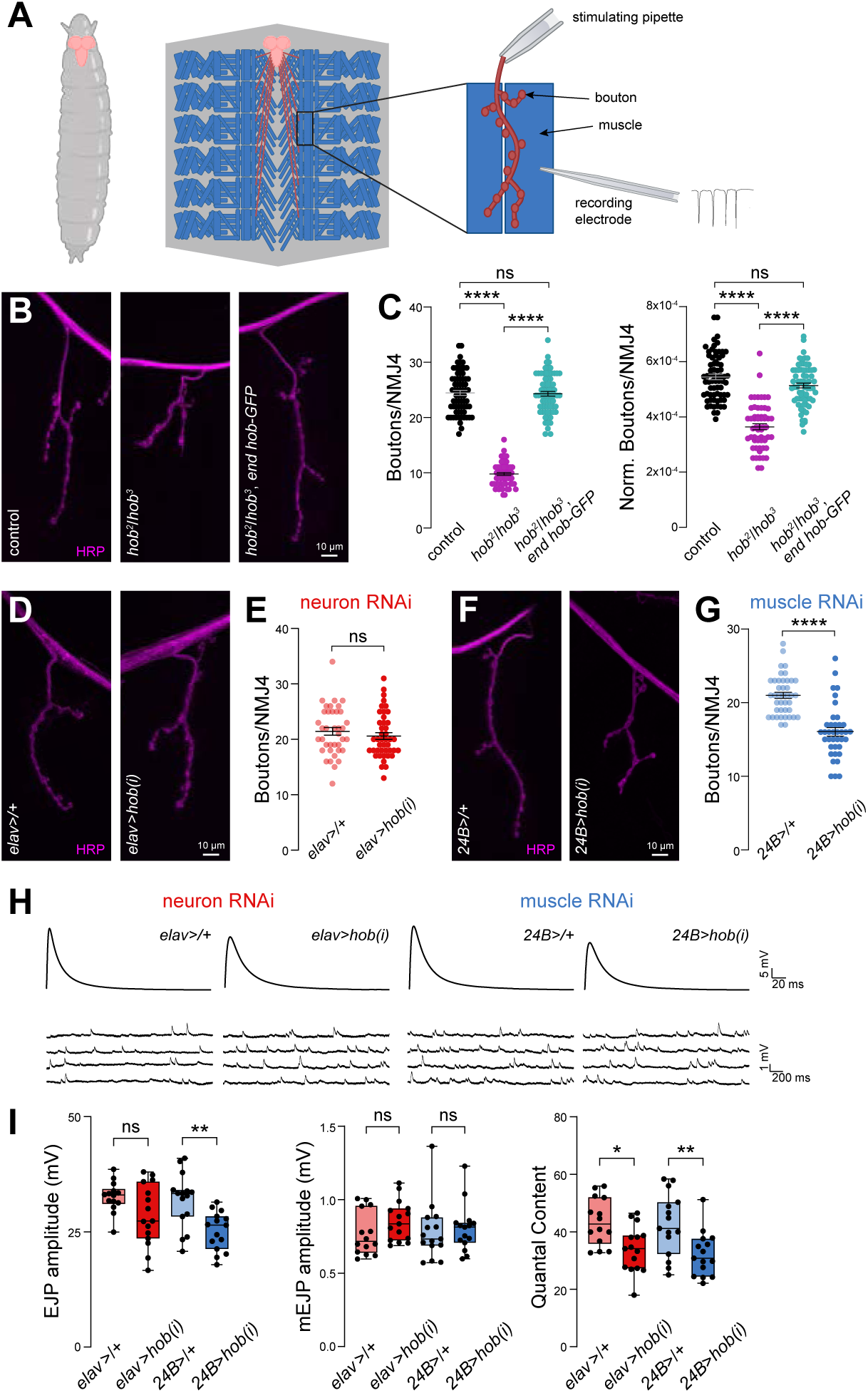
*hobbit* is required in the postsynapse to generate functional synapses. **(A)** Schematic of the brain/neurons (red) and the body wall muscles (blue) that comprise the larval neuromuscular junction (NMJ). The experimental paradigm for electrophysiology studies is shown on the right. **(B)** Representative images of NMJs in control, *hobbit* mutant (*hob^2^/hob^3^*), and rescue (*hob^2^/hob^3^, endogenous (end) hob-GFP*) larvae. **(C)** Bouton quantification in the same genotypes shown in panel B. Left graph shows total bouton number, right graph shows bouton number normalized to muscle area. Graphs shown mean ±S.E.M. with data points representing individual NMJs. *n*≥50 NMJs analyzed per genotype. Significance calculated by Kruskal-Wallis followed by Dunn’s multiple comparisons test. **(D)** Representative images of NMJs in control (*elav>/+*) and neuron-specific knockdown of *hobbit* (*elav>hob(i*)) larvae. **(E)** Bouton quantification in the same genotypes shown in panel D. Graphs shown mean ±S. E. M. with data points representing individual NMJs. *n*≥30 NMJs analyzed per genotype. Significance calculated by Mann-Whitney test. **(F)** Representative images of NMJs in control (*24B>/+*) and muscle-specific knockdown of *hobbit* (*24B>hob(i*)) larvae. **(G)** Bouton quantification in the same genotypes shown in panel F. Graphs shown mean ±S.E.M. with data points representing individual NMJs. *n*≥30 NMJs analyzed per genotype. Significance calculated by Mann-Whitney test. **( H-I )** Electrophysiological analysis of synapse function upon neuron-specific (red) or muscle-specific (blue) knockdown of *hobbit*. Representative traces for each genotype are shown in **H.** In **I**, left graph shows excitatory junction potentials (EJPs), middle graph shows miniature EJPs (mEJPs), and right graph shows quantal content (EJP amplitude/mEJP amplitude) with data points representing individual NMJs. Boxes extend from 25^th^ to 75^th^ percentiles; line in boxes indicates the median. Whiskers extend to the minimum and maximum values. *n*≥10 animals analyzed per genotype. Significance calculated by ordinary one-way ANOVA followed by Šídák’s multiple comparisons test for normally distributed data (EJP and QC) and by Kruskal-Wallis test followed by Dunn’s multiple comparisons test for non-normally distributed data (mEJP). *\*p*>0.05; \*\**p*<0.01; \*\*\*\**p*<0.0001; ns=not significant.

Communication between pre- and postsynaptic cells is essential to regulate synaptogenesis. For example, at the *Drosophila* NMJ, Wnt signals from presynaptic neurons and BMP signals from postsynaptic muscle are both required to coordinate synaptogenesis (61,62). Because the *hobbit* mutations analyzed in Fig. 4A-B are whole animal/systemic, we cannot make any conclusions about whether the protein is required pre- and/or postsynaptically to promote synaptogenesis. Thus, to determine if there was a specific pre- or postsynaptic role, we tested the impact of neuron- or muscle-specific knockdown of *hobbit*. Surprisingly, presynaptic (neuron-specific) knockdown had no effect on synaptic growth (Fig. 4D-E); however, postsynaptic (muscle-specific) knockdown significantly reduced bouton number (Fig. 4F-G). This result suggests that *hobbit* has a specific role in postsynaptic muscle to promote synaptogenesis.

We next used electrophysiological studies to assess NMJ function when *hobbit* is lost either pre- or postsynaptically. We measured spontaneous and evoked synaptic potentials through current clamp recordings and found that neurotransmission, as measured by evoked excitatory junction potentials (EJPs), was significantly reduced upon muscle-specific (but not neuron-specific) knockdown of *hobbit* (Fig. 4H-I). Muscle-specific knockdown of *hobbit* did not alter miniature EJP (mEJP) amplitude, which measures the response to spontaneous release of a single vesicle. Thus, the reduced EJP amplitude in muscle-specific loss of *hobbit* was due to a decrease in neurotransmitter release (quantal content; Fig 4I). This finding is consistent with the reduced number of boutons observed upon muscle-specific depletion of *hobbit* (see Fig. 4F-G). Neuronal knockdown of *hobbit* resulted in a small reduction in quantal content, reflecting the slight, non-significant decrease in EJP amplitudes and increase in mEJP amplitudes observed upon neuron-specific loss of *hobbit* (Fig. 4H-I). This result could suggest a mild neuronal deficit that worsens during development, contributing to the substantial locomotion defects observed in adult flies (see Fig. 3B). Together, these results show that *hobbit* plays an essential postsynaptic role in generating functional synapses during development.

## DISCUSSION

Mutations in BLTPs are associated with several neurodegenerative and neurodevelopmental disorders, including movement disorders seen in *VPS13A* disease and autism and schizophrenia associated with *BLTP2* mutations. Here, we used *Drosophila* to define both temporal and tissue-specific requirements for these BLTPs. Our data indicate that *VPS13A* function is required in neurons for maintenance of neuronal health and function during aging, while *BLTP2* is required in postsynaptic cells to promote synaptogenesis during development. These novel and independent functions for BLTPs provide new insights into the etiology of movement disorders and the physiological functions of bulk lipid transporters.

Beyond *VPS13A*, there are three other *VPS13* paralogs in the human genome; each is associated with a distinct neurodevelopmental or neurodegenerative disease: *VPS13B*-Cohen syndrome (63), *VPS13C*-early-onset Parkinson’s disease (64), *VPS13D*-spastic ataxia (65). Although all four paralogs share a common evolutionary origin (there is a single *Vps13* gene in the budding yeast *Saccharomyces cerevisiae*), this association with different diseases suggests that each paralog has evolved distinct functions. Since evidence so far suggests that BLTPs are non-selective for the lipids they transport, one conspicuous possibility is that specificity could be mediated by the subcellular localization of each VPS13 paralog. VPS13A primarily localizes to ER-lipid droplet and ER-mitochondria contacts (5), although it has also been reported at ER-PM contacts during specific cellular processes like erythrocyte differentiation (50,66). VPS13B localizes to the interfaces between Golgi cisternae (67), VPS13C localizes to ER-lipid droplet and ER-endosome/lysosome contacts (5,68), and VPS13D localizes to the Golgi and to ER-mitochondria and ER-peroxisome contacts (6). Although both VPS13A and VPS13D are detected at ER-mitochondria contacts, the pattern of localization is different and enrichment is driven by distinct adapter proteins (6). At ER-mitochondria contacts, non-vesicular lipid transfer of phospholipids like phosphatidylserine (PS), phosphatidic acid (PA), and phosphatidylcholine (PC) from their site of synthesis, the ER, to mitochondria is essential to maintain mitochondrial integrity and function (69). Given the importance of mitochondria in aging, myopathy, and neurodegeneration (70,71), the localization and function of VPS13A at ER-mitochondria contact sites is consistent with our findings that this protein is required for maintenance of neuronal and muscle health during aging. Moreover, this raises the possibility that the etiology of *VPS13*-related diseases is driven, at least in part, by the subcellular localization of each paralog.

Our data demonstrate that *BLTP2* is required in postsynaptic muscle to promote synaptogenesis at the NMJ. Although we do not yet know the mechanism by which *BLTP2* regulates synaptogenesis, we do know that *BLTP2* is required for protein secretion in several different cell types (34). Thus, *BLTP2/ hobbit* may be required for secretion of the BMP ligand *glass bottom boat* (*gbb*), which is released from postsynaptic muscle to promote synaptogenesis at the NMJ (62). This also raises the possibility that *BLTP2* could be required in all postsynaptic cells, including those in the central nervous system, particularly since neuron-specific knockdown of *hobbit* causes severe locomotion defects in adults. Alternatively, these deficits could reflect presynaptic roles hinted at by the small but significant reduction in evoked neurotransmitter release upon neuronal knockdown of *hobbit*. These phenotypes are consistent with the etiology of neurodevelopmental disorders, which hinge on the proper formation of functional synapses (33). Further study of the role of *BLTP2* in the central nervous system is likely to provide new insights into the role of this protein in neurodevelopmental disorders.

Given that BLTP-related diseases have primarily neurodevelopmental or neurodegenerative clinical manifestations, much of the focus has been on the function of these proteins in the nervous system. However, possible functions in other tissues should not be ignored. Our data suggests that myopathy is likely an independent comorbidity in *VPS13A* disease. Additionally, some patients with *VPS13A* disease present with abnormally shaped erythrocytes, called acanthocytes (1), suggesting that *VPS13A* likely has a role during erythrocyte formation. Indeed, one study suggests that VPS13A localizes to ER-PM contacts in differentiating erythrocytes, and this localization depends on the lipid scramblase *XK* (66). According to the Human Protein Atlas ((72); proteinatlas.org), none of the BLTP family members exhibit tissue-specific expression patterns, indicating that we should take a holistic approach to studying BLTP function as it relates to disease etiology. Additionally, although the membrane contact site localization of each BLTP has been identified, further study will be required to uncover the specific subcellular functions of BLTPs in human disease.

## ACKNOWLDEGEMENTS

Stocks obtained from the Bloomington *Drosophila* Stock Center (NIH P40OD018537) were used in this study. The authors thank the University of Wisconsin Translational Research Initiatives in Pathology laboratory (TRIP), supported by the UW Department of Pathology and Laboratory Medicine, UWCCC (P30 CA014520) and the Office of The Director-NIH (S10 OD023526) for use of its facilities and services. Diagrams in Figs. 1A, C, 2A, 3A, and 4A were created in https://BioRender.com.

## AUTHORS’ ROLES

Conceptualization: SDN, RST, KMOG, AB; Methodology: SDN, RST, KOG, AB; Validation: SDN, RST, SJG, KMOG, AB; Investigation: SDN, RST, SJG; Data curation: SDN, RST, SJG; Writing - original draft: SDN.; Writing - review & editing: SDN, RST, SJG, KMOG, AB; Visualization: SDN, AB; Supervision: KMOG, AB; Project administration: KMOG, AB; Funding acquisition: KMOG, AB.

## SUPPLEMENTARY DATA

**Figure S1.**
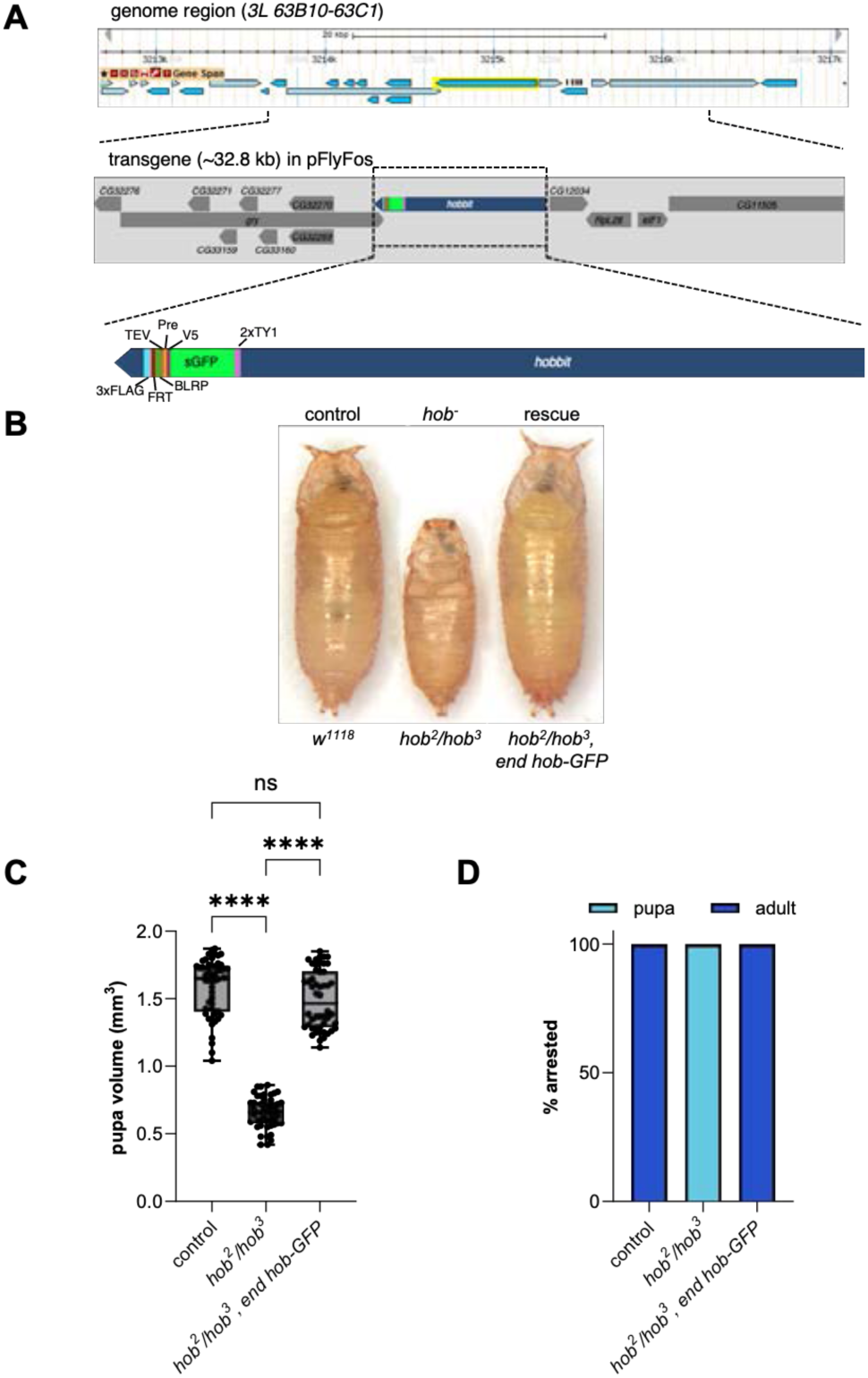
Generation and validation of *endogenous hobbit-sGFP*. **(A)** Schematic of the transgene containing *endogenous hobbit-GFP*. We obtained a fosmid containing ∼32.8kb of chromosome *3L*, including *hobbit* tagged at the C-terminus with superfolder GFP (sGFP) and other epitope tags (40). Sanger sequencing was used to confirm that the endogenous *hobbit* stop codon was deleted, and that the sGFP and other tags were present and in the correct reading frame. We then generated transgenic flies containing this construct inserted on chromosome *3R* at *89E11*. (**B-D**) Rescue of *hobbit* mutant animals with *end hob-GFP*. One copy of *end hob-GFP* is sufficient to rescue the small body size and lethality of *hob* mutants, seen by representative pupa pictures in **(B)**, quantification of body size in **(C)**, and lethal phase analysis in **(D)**. In panel C, boxes extend from 25^th^ to 75^th^ percentiles; line in boxes indicates the median. Whiskers extend to the minimum and maximum values. Pupae in panel B were initially captured in the same image but were individually rotated and aligned post-acquisition to improve image aesthetics. *n*=50 animals per genotype. Significance calculated by Kruskal-Wallis followed by Dunn’s multiple comparison test. \*\*\*\**p*<0.0001; ns=not significant.

## SUPPLEMENTARY EXTENDED METHODS

### Adult climbing and larval crawling

For adult climbing, groups of *n*∼10 flies were collected at 1-2 days old and allowed to age for the appropriate time. Then, each group of flies of the appropriate age and genotype was placed into a 50 mL graduated cylinder and allowed to acclimate for 3 min. All flies were then tapped to the bottom of the cylinder, and the number of flies climbing 8 cm in 12 s was counted. This procedure was repeated three times for each group of flies, and the average of the three trials is reported on the graphs. Three independent biological replicates with approximately equal numbers of male and female flies were analyzed for each genotype. Note that the same *elav>luc*, *elav>Vps13(i)*, *24B>luc*, and *24B>Vps13(i)* flies were used for climbing assays during aging and for lifespan analysis. During these experiments, flies were transferred to fresh food several times weekly, and the number of living flies was recorded prior to obtaining climbing data weekly. For larval crawling, third-instar larvae were washed with distilled water and placed in the center of a 150 mm petri dish filled with 1% agarose made with blue ink for contrast. Groups of five larvae were allowed to acclimate for 1 min. Larvae were then recorded for 75 s using a Sony FDR-AX53 4K Handycam (30 frames per second). Total distance crawled was calculated using the wrMTrckr plugin (42) in FIJI (43,44).

### Larval neuromuscular junction (NMJ) imaging and synaptic bouton quantification

Third-instar larvae were dissected in Ca^2+^-free saline and fixed for 6 min in Bouin’s fixative (Sigma Aldrich, HT10132). Dissected larvae were washed and permeabilized in PBS with 0.1% Triton X, blocked in PBS containing 0.1% Triton X and 1% BSA and 5% NGS for 30 min at room temperature (RT) or overnight at 4°C, followed by incubation in primary antibodies overnight at 4°C and secondary antibodies for 4 h at room temperature, then mounted in Vectashield (Vector Laboratories, H-1000-10). The following antibody was used at the indicated concentration: Alexa Fluor 647-contugated anti-HRP at 1:500 (Jackson ImmunoResearch Laboratories, Inc, 123-605-021; RRID: AB_2338967).

### Histology and neurodegeneration index quantification

Fly heads were severed and fixed overnight at 4°C in a 6:3:1 solution of ethanol:chloroform:glacial acetic acid, then washed and stored in 70% ethanol for histology. Heads were processed into paraffin using standard histological procedures; serial 5 µm sections spanning the whole fly head were obtained and stained with hematoxylin and eosin (H&E) (UW-Madison Translational Research Initiatives in Pathology lab (TRIP)).

### Electrophysiology

Third-instar male larvae were dissected in low Ca^2+^ modified hemolymph-like saline (HL3: 70 mM NaCl, 5 mM KCl, 15 mM MgCl2, 10 mM NaHCO3, 115 mM sucrose, 5 mM trehalose, 5 mM HEPES, and 0.2 mM Ca^2+^, pH 7.2). Recordings were performed at muscle 6 of abdominal segments A3 and A4 in HL3 containing 0.6 mM Ca^2+^. Recordings were performed on a Nikon FN1 microscope using a 40× (0.80

NA) water-dipping objective and acquired using an AxoClamp 900A amplifier, Axon Digidata 1550B low-noise data acquisition system, and pClamp 11.0.3 software (Molecular Devices). Electrophysiological sweeps were digitized at 10 kHz and filtered at 0.1 kHz. Excitatory junction potentials were recorded using 1.0 mm × 0.58 mm borosilicate glass sharp electrodes (10 and 15 MΩ resistance) filled with 3 M KCl. Miniature excitatory junctional potentials (mEJPs) were recorded with no external stimulation. For each recording, at least 100 mEJPs were analyzed using Mini Analysis (Synaptosoft) to obtain a mean mEJP amplitude value per muscle. A 1.5 mm × 1.12 mm borosilicate polished glass electrode was used to suction cut motor axons, which were stimulated at 0.2 Hz with 0.5 ms pulses using an A-M Systems isolated pulse stimulator 2100 to elicit EJPs. Stimulus intensity was adjusted to consistently elicit compound responses from both type Ib and Is motor neurons. Recordings were only performed if the resting membrane potential was between 50 and 80 mV and muscle input resistance was 4 MΩ or higher.

### Statistical analyses

Quantification was conducted blind to genotype. Sample sizes were based on prior published studies. Statistical analyses were conducted in GraphPad Prism 10. Normality was determined by the D’Agostino–Pearson omnibus test. The Mann–Whitney U test was used for single comparisons of non-normally distributed data. For single comparisons of means, unpaired *t*-tests with Welch’s correction were used. For multiple comparisons of normally distributed data, we performed ANOVA followed by Šídák’s test. For multiple comparisons of non-normally distributed data, we performed a Kruskal-Wallis test followed by Dunn’s test. Reported values are mean ± SEM or SD as indicated. Unless otherwise indicated, significance refers to the indicated genotype compared with control. *p* values, statistical test used, and sample sizes are reported in each figure legend.

